# Cryo-EM structure of the potassium-chloride cotransporter KCC4 in lipid nanodiscs

**DOI:** 10.1101/805267

**Authors:** Michelle S. Reid, David M. Kern, Stephen G. Brohawn

## Abstract

Cation-chloride-cotransporters (CCCs) catalyze transport of Cl^−^ with K^+^ and/or Na^+^ across cellular membranes. CCCs play roles in volume regulation, neural development and function, audition, blood pressure regulation, and renal function. CCCs are targets of drugs including loop diuretics and their disruption is implicated in pathophysiologies including epilepsy, hearing loss, and the genetic disorders Andermann, Gitelman, and Bartter syndromes. Here we present the cryo-EM structure of a CCC, the *Mus musculus* K^+^-Cl^−^ cotransporter KCC4, in lipid nanodiscs. The structure, captured in an inside-open conformation, reveals the architecture of KCCs including an extracellular domain poised to regulate transport activity through an outer gate. We further identify substrate K^+^ and Cl^−^ binding sites and provide a structural explanation for differences in substrate specificity and transport ratio between CCCs. These results provide mechanistic insight into the function and regulation of a physiologically important transporter family.

## Introduction

CCCs in mammals include the potassium-chloride cotransporters KCC1-4, the sodium-potassium-chloride cotransporters NKCC1-2, the sodium-chloride cotransporter NCC, and CCC8-9 (Figure S1)^1–3^. First characterized as modulators of red blood cell volume^4, 5^, CCCs are now appreciated to play critical roles in cellular volume regulation, modulation of neuronal excitability, renal function, auditory system function, transepithelial transport, and blood pressure regulation^2, 3, 6^. CCCs are targets of drugs including the thiazide and loop diuretics hydrochlorothiazide, furosemide, and bumetanide and their disruption is associated with congenital hydrocephaly, epilepsy, hearing loss, Andermann syndrome, Gitelman syndrome, and Bartter syndrome^3, 7, 8^.

KCCs are important for K^+^ and Cl^−^ homeostasis, including in establishing low neuronal cytoplasmic Cl^−^ concentrations critical for inhibitory neurotransmission, and in volume regulation in many cell types^2, 9–11^. Among KCCs, KCC4 is most strongly activated by cell swelling and high internal [Cl^−^] and is uniquely active in acidic external environments^2, 9^. KCC4 is expressed in tissues including the heart, nervous system, kidney, and inner ear and mice lacking KCC4 display progressive deafness and renal tubular acidosis^2, 10–12^. Hearing loss in these animals is due to disrupted K^+^ recycling by Dieter’s cells in the cochlea and hair cell excitotoxicity, while renal tubular acidosis is due to impaired Cl^−^ recycling by α-intercalated cells in the kidney distal nephron^12^.

CCCs display varied substrate specificity and transport stoichiometry despite sharing a common amino acid-polyamine-organocation (APC) superfamily fold^13–15^. KCCs cotransport K^+^:Cl^−^ in a 1:1 ratio, NKCCs cotransport 1K^+^:1Na^+^:2Cl^−^, and NCCs cotransports 1Na^+^:1Cl^−^ ^2, 3^. One consequence of this difference is that under typical conditions (with [K^+^]_in_:[K^+^]_out_ > [Cl^−^]_out_:[Cl^−^]_in_), transport by KCCs is outwardly directed while transport by NKCCs/NCCs is directed into cells^2, 3^. CCCs have two distinctive elaborations on the APC fold. First, the scaffold is followed by a C-terminal domain (CTD) important for regulating expression, trafficking, and activity including through phosphorylation or dephosphorylation of CTD sites in response to cell swelling^16–19^. Second, CCCs contain a “long extracellular loop” with predicted disulfide bonds and glycosylation sites that differs in position and structure between CCCs; it is formed by the region between TM5-TM6 in KCCs and between TM7-TM8 in NKCCs^14, 20^.

KCCs are present as monomers and dimers in cells and modulation of quaternary state has been implicated in transporter regulation. A shift from monomeric to dimeric KCC2 during development coincides with an increase in transport activity that results in chloride extrusion from neurons (the excitatory-to-inhibitory GABA switch)^21–23^. Homodimerization is likely largely mediated through CTD interactions, as observed in the recent cryo-EM structure of NKCC1^21^ and calpain-mediated proteolysis of the KCC2 CTD is associated with a decrease in transporter activity^23^. In addition to self-associating, KCCs heterodimerize with other CCCs and interact with other membrane proteins including ion channels^22, 24^.

Here we report the structure of *Mus musculus* KCC4 in lipid nanodiscs determined by cryo-EM. The structure reveals unique features of KCCs and, together with functional characterization of structure-based mutants, provides insight into the basis for ion substrate selectivity, transport stoichiometry, and regulation of activity in potassium-chloride cotransporters.

## Results

### Structure of KCC4 in lipid nanodiscs

*Mus musculus* KCC4 was heterologously expressed in *Spodoptera frugiperda* (Sf9) insect cells for purification and structure determination (Figure S2,S3). To assess the activity of KCC4 in these cells, we utilized an assay that depends on the ability of KCCs to transport Tl^+^ in addition to K^+ 25^. In cells loaded with the Tl^+^-sensitive fluorophore FluxOR red, Tl^+^ uptake from the extracellular solution results in an increase in fluorescence signal (Figure 1A). Cells infected with virus encoding KCC4, but not cells infected with a virus encoding an anion-selective volume-regulated ion channel SWELL1^26^ or uninfected Sf9 cells, displayed increased fluorescence over time consistent with KCC4 activity (Figure 1B,C). No significant difference in activity was observed between N- and C-terminally GFP-tagged mouse KCC4 (Figure 1B,C), in contrast to a previous report for KCC2^27^, and C-terminally tagged KCC4 was used for subsequent study.

**Figure 1.**
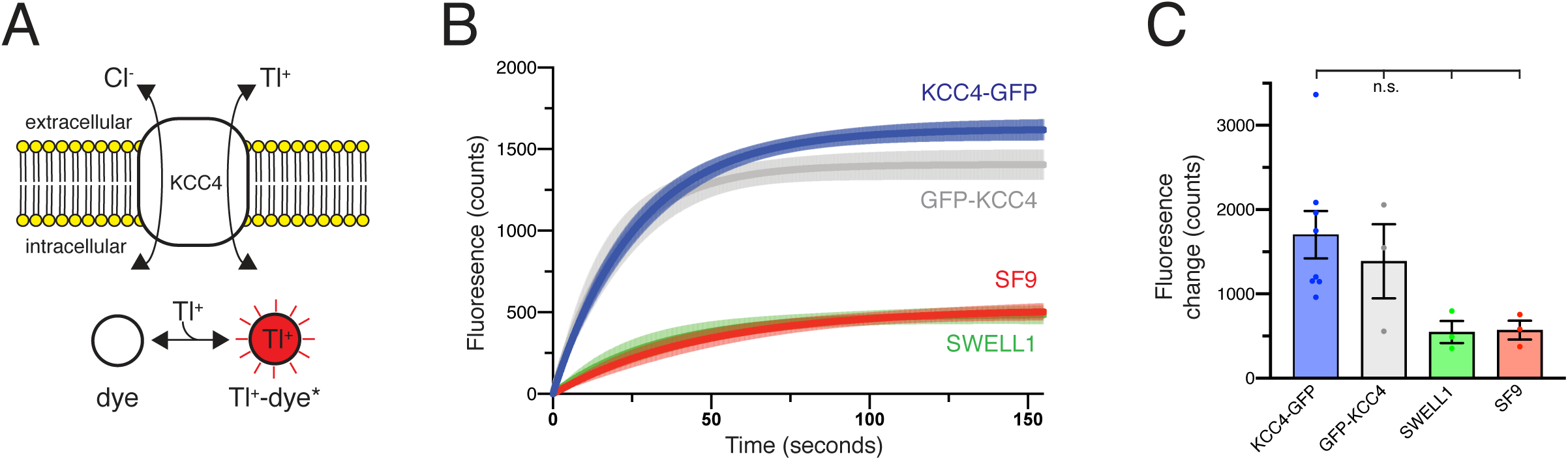
Transport activity of mouse KCC4. (A) A Tl+ uptake assay for KCC4 activity. KCC4 activity in SF9 cells results in Tl+ uptake and increased fluorescence of the Tl+ sensitive dye FluxOR Red. (B) Fluorescence values as a function of time for each construct assayed. Lines are global exponential fits to all data with 95% confidence intervals shown for KCC4-GFP (n=8, blue), GFP-KCC4 (n=3, gray), SWELL1 (n=3, green), and uninfected SF9 cells (n=3 red). (C) Quantification of experiments shown in (B). Average final fluorescence over 25s is plotted. KCC4-GFP 1703±280.1 (n=8) GFP-KCC4 1387±440.0 (n=3) SWELL1 546.6±131.3 (n=3) SF9 570.2±110.6 (n=3). Statistical differences assessed with one-way Anova (n.s. = not significant, * = p<0.05).

We reconstituted KCC4 into lipid nanodiscs in order to study the structure of the transporter in a native-like membrane environment. KCC4 was extracted, purified in detergent, and exchanged into nanodiscs formed by the membrane scaffold protein MSP1D1 and a mixture of phospholipids that approximates their representation in neuronal membranes (2:1:1 molar ratio DOPE:POPC:POPS (2-dioleoyl-sn-glycero-3-phosphoethanolamine:1-palmitoyl-2-oleoyl-sn-glycero-3-phosphocholine:1-palmitoyl-2-oleoyl-sn-glycero-3-phospho-L-serine)) (Figure S2)^28, 29^. KCC4-MSP1D1 particles are similar in size and shape to KCC4 particles in detergent micelles by cryo-EM, but show improved distribution in thin ice which facilitated reconstruction to high resolution (Figure S4).

An unmasked reconstruction of KCC4 in nanodiscs shown in Figure 2A is contoured to highlight the position of the lipid membrane surrounding the transmembrane region of the transporter. To achieve the highest resolution reconstruction, the nanodisc density was subtracted and particles were subjected to focused classification and subsequent refinement (Figure S5). The resulting map, at 3.6 Å overall resolution, enabled complete de novo modeling of the transmembrane and extracellular region of KCC4 and includes two partial extracellular glycosylation sites, a bound K^+^ ion, and a bound Cl^−^ ion (Figures 1C,1D,S6, and S7).

**Figure 2.**
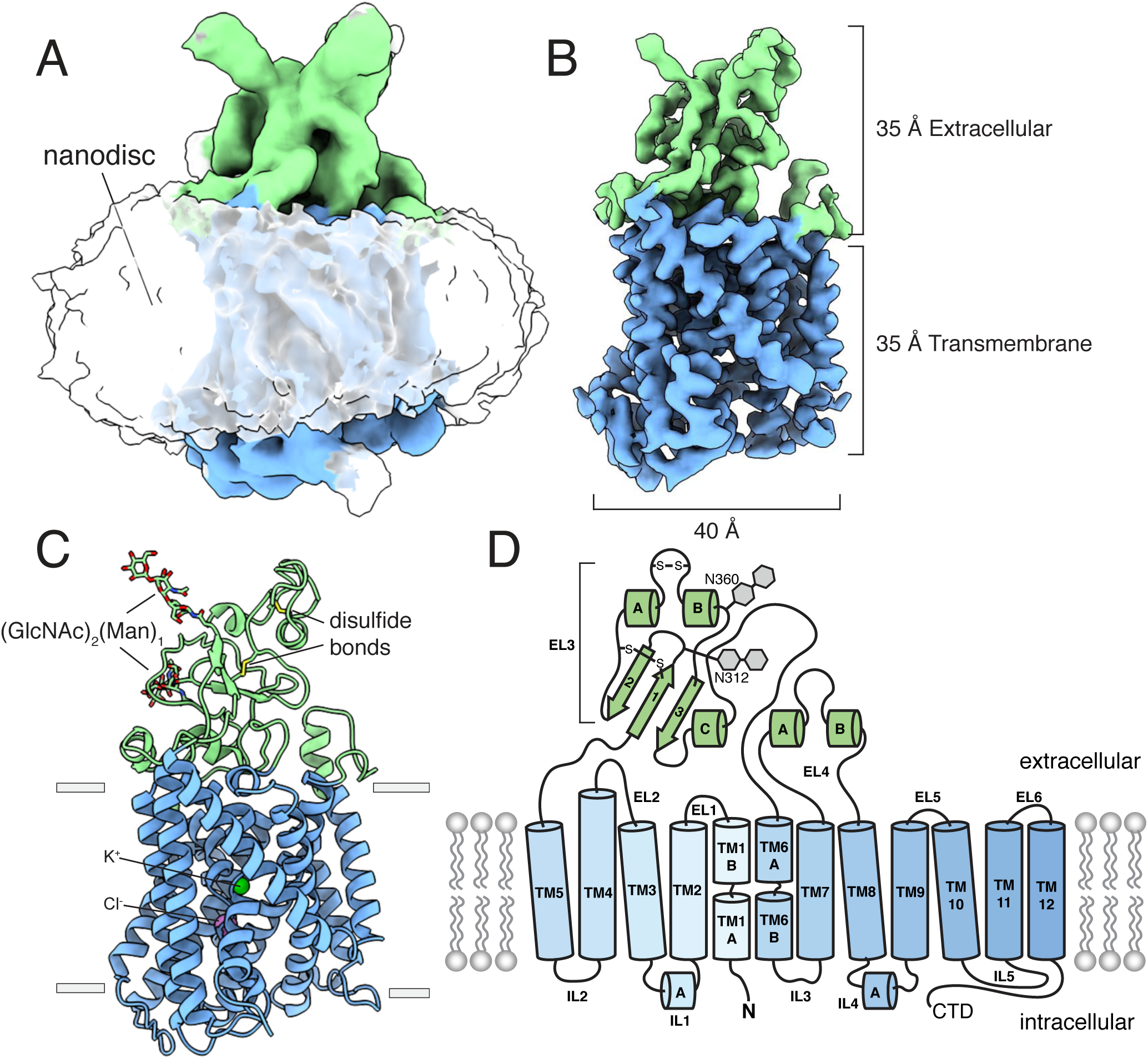
Structure of mouse KCC4 in lipid nanodiscs. (A) Cryo-EM map from an unmasked refinement viewed from the membrane plane showing the position of nanodisc, transmembrane region (blue), and extracellular region (green). (B) Final map, (C) corresponding atomic model, and (D) cartoon representation of KCC4. In (C), bound K+ and Cl− ions are shown as green and violet spheres, respectively. Three disulfides and two N-linked glycosylation sites are shown as sticks and labeled in the cartoon.

### Overall architecture

KCC4 is monomeric in the nanodisc structure, in contrast to the recent homodimeric structure of *Danio rerio* NKCC1^30^. Density for the N-terminal region and C-terminal domain (CTD), which together comprise approximately half the expressed protein mass, is not observed in the cryo-EM maps (Figure 2, S1, and S2). The N-terminal region is weakly conserved, variable in length among CCCs (Figure S1), and is likewise unresolved in the structure of NKCC1^30^. We presume it is highly flexible in KCC4. The C-terminal domain, however, is well conserved, has documented roles in regulation, expression, and trafficking^1, 2,13,14,16^, and mediates homodimerization of NKCC1 and the Archaean CCC (MaCCC)^30, 31^.

We evaluated possible explanations for the difference in oligomeric state between KCC4 and NKCC1. Proteolytic cleavage of the dimerization-mediating CTD was excluded because the CTD was identified by mass spectrometry of purified KCC4 with high coverage (47%) and abundance (81% of all KCC4 peptides) (Figure S3). Disruption of putative homodimers during purification or sample preparation was excluded for the following reasons: (i) The portion of KCC4 in an early-eluting broad peak from a sizing column (Figure S2A) displays nonspecific aggregation by cryo-EM. (ii) KCC4 is monomeric before and after reconstitution in nanodiscs as assessed by cryo-EM (Figures S2,S4). (iii) Cross-linking of purified KCC4 was observed only at high concentrations of crosslinker (Figure S2F) and was reduced when KCC4 was first deglycosylated (Figure S2E,F), suggesting some cross-linking in glycosylated KCC4 is from intermolecular glycan-glycan or protein-glycan linkages rather than through transmembrane regions or CTDs^21^. We conclude that the CTD is flexibly attached to the transmembrane region. Consistently, we observe a progressive loss of detailed features and a decrease in local resolution in TM11 and TM12 that connect the CTD to the core transmembrane region (Figure S7B, S8). Some two-dimensional class averages show a blurred cytoplasmic feature around the position we expect the CTD to emerge (Figure S4), although attempts to classify distinct conformations of this feature were unsuccessful. We conclude that the monomeric structure reported here corresponds to full-length mouse KCC4 with intrinsically flexible and/or disordered terminal regions.

### Transporter conformation

KCC4 adopts an inward-open conformation. The outer surface of the transporter is sealed from the extracellular solution, while a continuous cavity extends from the center of the transmembrane region to the cytoplasmic side (Figure 3). The transmembrane region consists of twelve helices (TM1-TM12) with TM1-TM5 related to TM6-TM10 through an inverted repeat. TM2 and TM11 in KCC4 are linked by a membrane buried disulfide bond between amino acids C163 (TM2) and C626 (TM11) that are conserved among KCCs (Figure S1).

**Figure 3.**
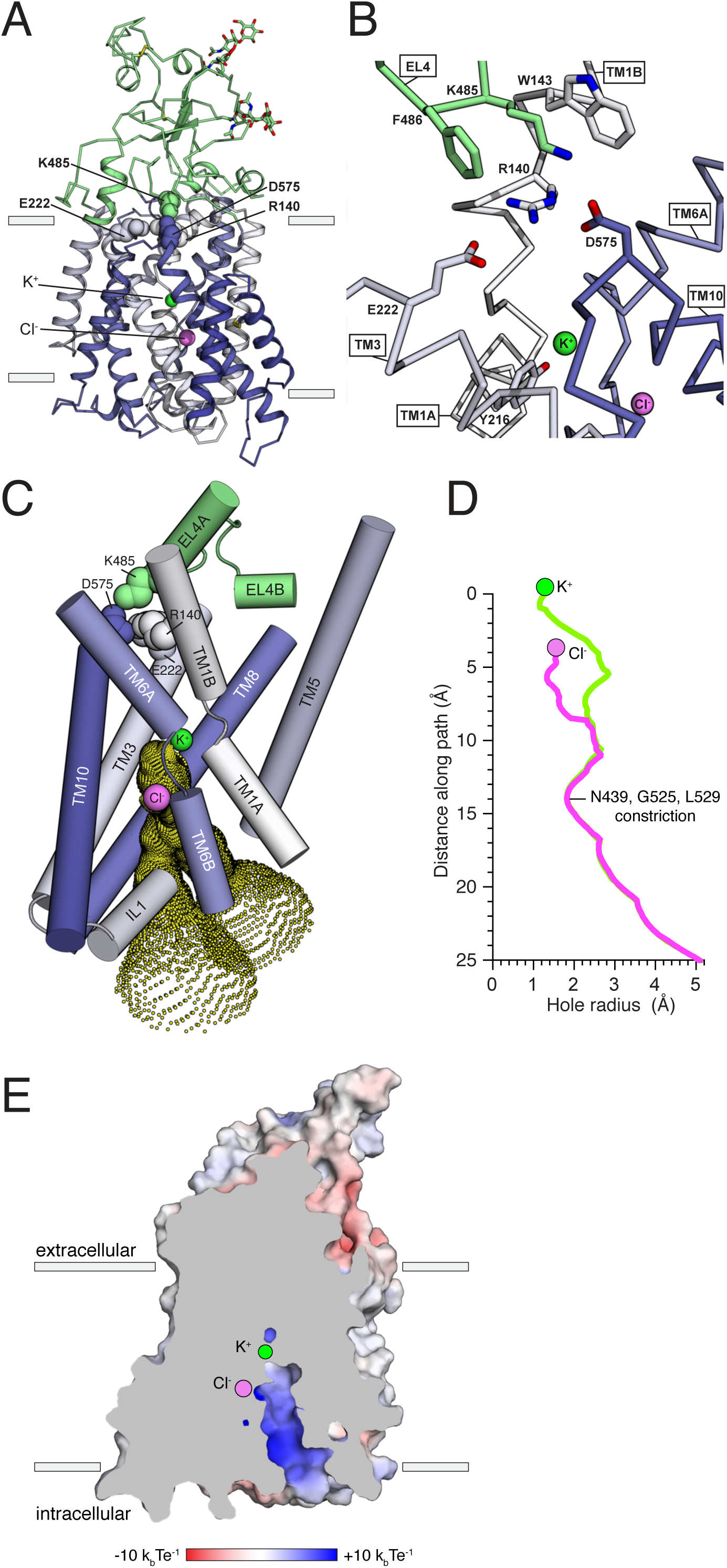
Inward-open conformation of KCC4. (A) Cartoon representation of KCC4 colored with extracellular region green and transmembrane region colored in a gradient from white to blue from N- to C-terminus. Ions and residues forming an extracellular gate are shown as spheres. (B) Close-up view of extracellular gate. Residues forming the electrostatic interaction network are shown as sticks. (C) View of the open pathway to the intracellular ion binding sites. Helices surrounding the ion binding sites are shown as cylinders. Yellow dots demarcate the surface of a bifurcated tunnel that connects the ion binding sites to the cytoplasmic solution. (D) Radius of the ion access tunnel as a function of distance along the path for Cl− (pink) and K+ (green). (E) Electrostatic surface representation of KCC4 sliced to show one leg of the cytoplasmic access tunnel.

A prominent feature of KCC4 is a large extracellular domain, unique among proteins of known structure, extending ∼35 Å above the membrane. It is formed by EL3 (the long extracellular loop^20^) and EL4, which pack together and cover approximately two-thirds of the transporter outer surface (Figure 2C-D, 3A). Sequence comparison suggests this domain is conserved in all KCCs and is not found in other CCCs (Figure S1). The structure consists of a short three-stranded antiparallel beta sheet (EL4 S1-3), five short helices (EL3 HA-C and EL4 HA,B), and regions without regular secondary structure. It is stabilized by two disulfide bonds (C308-C323 and C343-C352)^20^ and decorated with N-linked glycosylation sites^2, 32^ conserved among KCCs. Non-protein density consistent with glycosylation is present at four previously identified sites (N312, N331, N344, and N360)^32^ and we model partial carbohydrate chains at the two stronger sites (N312 and N360). Notably, the carbohydrate chain at N312 projects from the EH3 S1-S2 loop underneath an extended segment that leads to TM6A. This arrangement may stabilize the extracellular domain and couple it to movements in TM6A, which moves between functional states in other APC transporters^15, 33^. This structural role for glycosylation may explain the functional defects associated with non-glycosylated mutants of KCC4^32^.

The position of the extracellular domain suggests its involvement in conformational changes during the KCC transport cycle. A segment of the extracellular domain close to the membrane forms part of the constriction that seals the internal vestibule from the extracellular solution. This is likely the extracellular gate based on comparison to other APC transporters (Figure 3A,B)^15^. In KCC4, residues in EL4, TM1, TM3, and TM10 form an electrostatic and hydrophobic interaction network that seals the gate (Figure 3A,B). R140 on TM1B extends towards the extracellular solution to interact with D575 on TM10 and E222 on TM3. The extracellular domain is positioned immediately above through an interaction between K485 on EL4 and D575. The outer portion of TM1B contributes W143 which, together with F486, surrounds K485 as it projects towards TM10. This is reminiscent of the extracellular gate in LeuT formed by an electrostatic interaction between TM1 and TM10 (R30 and D404) and capped by EL4 through an interaction with TM10 (D401 and A319)^34^. By analogy, opening of the KCC4 extracellular gate is likely to require “unzipping” the electrostatic network and rotation of EL4 and the extracellular domain away from the surface of the transmembrane region^34, 35^.

On the intracellular side of KCC4, a hydrophilic cavity is formed by TM1, TM3, TM6, and TM8 that exposes the inside of the transporter to the cytoplasm (Figure 3C-E). At the top of this cavity are Cl^−^ and K^+^ binding sites. The cavity forms a bifurcated pathway for ion access to these sites, splitting into two routes approximately halfway through the tunnel due to the position of side chains of N439, R440, and R528. Both sides are open to an essentially equivalent degree (Figure 3D). The only constriction outside of the local area surrounding the ions is formed at the position of N439 (from TM6B), G525, and L529 (from TM8) where the cavity narrows to ∼3.6 Å in diameter, which is still sufficiently large for passage of K^+^ and Cl^−^ ions. Within ∼3 Å of each ion, the cavity narrows such that it would require at least partial ion dehydration.

The cavity surface is markedly electropositive (Figure 3E). From the intracellular solution up to the position of the Cl^−^, charged and polar side chains (from R440, R528, R535, N131, N274, N439, and N521), backbone amides (from IL1), and a helical dipole (from TM6B) contribute electropositive character. Since intracellular Cl^−^ ions are typically present at lower concentrations than K^+^ ions, this may serve to favor accumulation of the less abundant substrate near its binding site within transporter. Above the Cl^−^ site and around the K^+^ site, the accessible surface becomes electronegative and would favor cation binding. The extracellular surface of the transporter outside of the sealed gate is markedly electronegative. How this relates to ion binding or release in outward-open states awaits additional structural information.

### Substrate ion binding sites

The central discontinuities in TM1 and TM6 result in protein backbone carbonyls and amides not involved in regular hydrogen bonding that are utilized for substrate binding in other APC transporters^15^. Around this region, we observe two prominent non-protein density features (Figure 4 A,D). Based on structural, functional, and comparative analyses described below, we model these sites as bound K^+^ and Cl^−^ ions.

**Figure 4.**
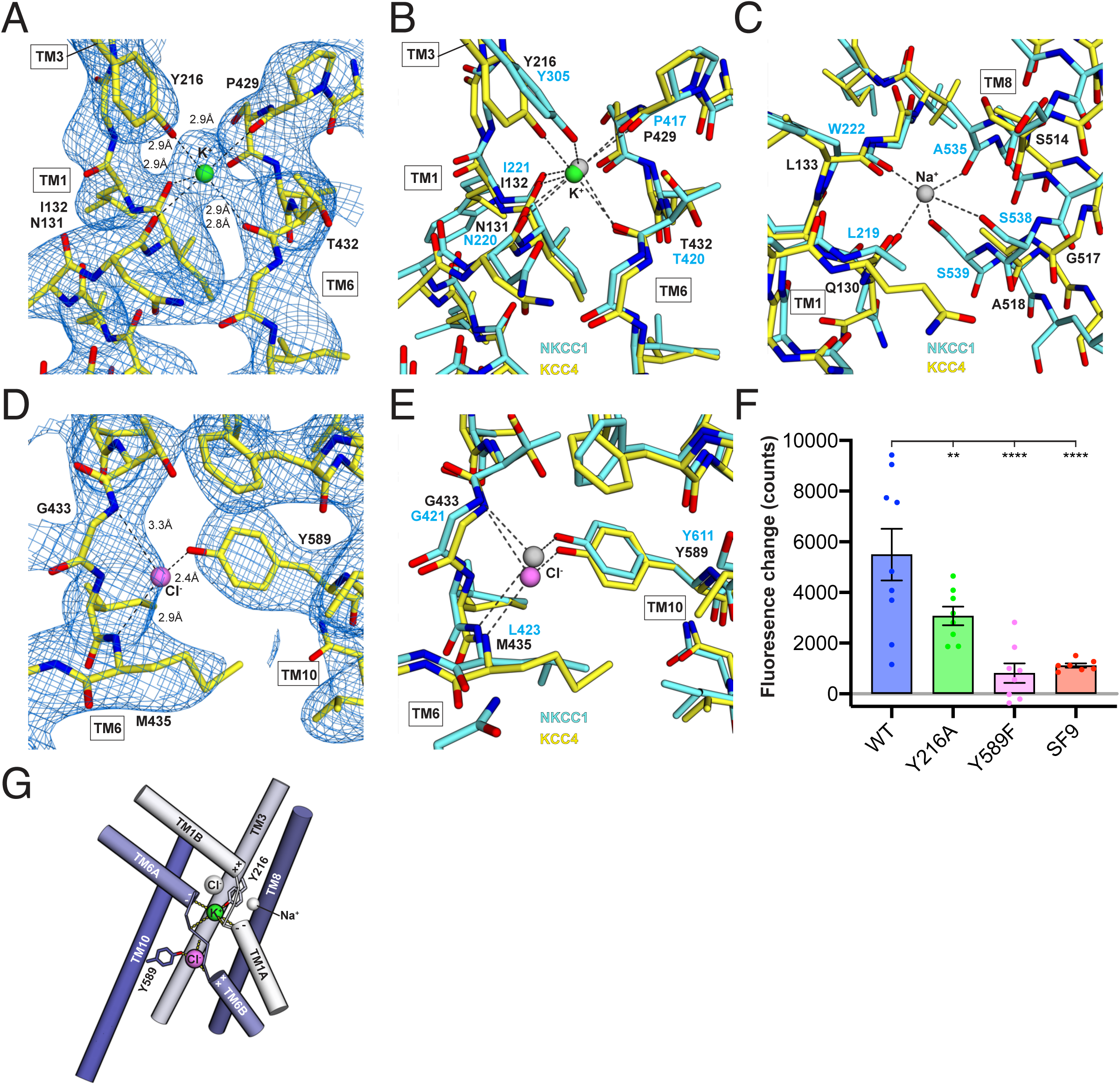
Ion binding sites. (A) K+ binding site. The cryo-EM map is shown as blue mesh and KCC4 is colored with carbon yellow, oxygen red, nitrogen blue, and K+ green. K+-coordination environment is indicated with dashed lines. (B) Superposition of K+ binding sites in KCC4 (depicted as in (A)) and NKCC1 (PDB:6NPL) colored with carbons cyan and K+ gray. (C) Superposition of Na+ binding site Danio rerio NKCC1 and analogous region in KCC4. The position of the Na+ ion is inferred from SiaT (PDB:5NVA). (D) Cl− binding site and (E) superposition of Cl− binding site in KCC4 with analogous site in NKCC1. (F) Activity of mutants in ion binding sites. Average final fluorescence over 25s is plotted. Wild-type KCC4 5496±1018 (mean±SEM, n=9), Y216A 3073±365.6 (n=8), Y589F 818.5±385.5 (n=8), and uninfected SF9 1116±82.64 (n=7). Statistical differences assessed with one-way Anova (** p<0.01, ****p<0.0001). (G) Model for substrate binding and transport stoichiometry in CCC transporters. Helices are shown as cylinders with ion coordination in KCC4 shown as dashed lines to green K+ and violet Cl−. Helix dipoles in discontinuous helices TM1 and TM6 are indicated. Additional ion binding sites in NKCC1 not observed in KCC4 (an upper Cl− and a Na+ site) are shown as transparent spheres.

The stronger of the two densities between TM1, TM6, and TM3 is modeled as a K^+^ ion. It is surrounded by electronegative groups contributed by backbone carbonyls (N131 and I132 in TM1 and P429 and T432 in TM6) and a tyrosine hydroxyl from Y216 in TM3. The distances between electronegative groups and the ion are consistent with K^+^ binding (2.8 – 2.9 Å). The electronegative helix dipoles created by TM1A and TM6A may additionally contribute to a favorable electrostatic environment for cation binding. The coordinating tyrosine is conserved in all CCC family members that transport K^+^. In NCC, the position corresponding to Y216 is substituted by a histidine, which likely explains its K^+^-independence (Figure S1).

The second site, between TM6 and TM10, is modeled as a Cl^−^ ion. It is surrounded by electropositive groups from backbone amides (G433 and I434 in TM6) and a tyrosine hydroxyl from Y589 in TM10. The electropositive helix dipoles created by TM1B and TM6B may additionally stabilize anion binding. The interaction distances and coordination environment are reminiscent of Cl^−^ sites in CLC transporters^36^ and the coordinating tyrosine is conserved across CCCs.

To validate the assignment of the K^+^ and Cl^−^ sites and test the importance of coordinating residues in transporter activity, we mutated Y216 and Y589 that contribute to the binding sites and assessed transporter activity. Mutations at both sites resulted in a marked reduction in transport activity in the Tl^+^-flux assay: mutation of the K^+^-coordinating Y216 to alanine resulted in a 44% reduction in activity and mutation of the Cl^−^-coordinating Y589 to phenylalanine resulted in an 85% reduction in activity compared to wild-type KCC4. We conclude these two sites are critical for KCC4 activity.

How does substrate binding in the 1:1 K^+^:Cl^−^ cotransporter KCC4 differ from the 1:1:2 Na^+^:K^+^:Cl^−^ cotransporter NKCC1^30, 37^? Overlays of the relevant sites are shown in Figure 4B,C,E. The KCC4 K^+^ and Cl^−^ sites correspond closely to sites for the same ions in NKCC1. However, the proposed Na^+^ site in NKCC1 is dramatically reorganized in KCC4. In KCC4, TM8 is rotated farther away from TM1 and two consecutive Na^+^-coordinating serines in NKCC1 (conserved in all Na^+^-transporting CCCs) are substituted by glycine and alanine in KCC4 (and in KCC1-3) (Figure S1). The consequence is a loss of three of the five Na^+^-coordinating positions, providing a structural explanation for Na^+^-independence in KCCs. A second Cl^−^ site in NKCCs extracellular to the K^+^ site is structurally conserved in KCC4, but we do not observe evidence for ion occupancy at this site in the structure, suggesting it may be lost in KCC4 in conjunction with the Na^+^-site (Figure 4G).

## Discussion

The structure of monomeric KCC4 in lipid nanodiscs is consistent with evidence for KCC monomers in cells in addition to homodimers and other oligomers^22, 23, 38^. Comparison of residues in KCC4 that correspond to those in the NKCC1 homodimerization interface^30^ suggest KCC4 could similarly self-interact. An intriguing hypothesis is that monomeric and dimeric KCC4 are functionally distinct and modulated differently; monomers may function independently of flexibly attached CTDs, while dimerization, which involves close juxtaposition of CTDs and transmembrane regions in NKCC1, could enable regulation of transporter activity through CTD posttranslational modifications^14, 22, 23, 38^. Consistent with this notion, a monomeric to dimeric transition in KCC2 has been correlated with an increase in transporter activity^22^ and can be regulated by phosphorylation of the CTD^38^, while proteolytic cleavage of the KCC2 CTD correlates with a decrease in activity^23^. Alternatively, the ability of KCC4 to hetero-oligomerize with other CCCs and other membrane proteins may be associated with weaker self-interaction^24, 39–41^.

The structure of KCC4 provides insight into the architecture of KCCs and the mechanistic basis for coupled K^+^:Cl^−^ transport. KCC4 is observed in an inside-open conformation that exposes the inside of the transporter to the cytoplasm through a wide and electropositive tunnel that may serve to concentrate the less abundant intracellular substrate Cl^−^. This conformation is similar to that observed in NKCC1 and presumably represents the lowest energy state for both CCCs in symmetrical salt concentrations in the absence of a transmembrane electrical gradient^30^. We identify K^+^ and Cl^−^ ions around central discontinuities in TM1 and TM6. A Na^+^ site in the 1:1:2 Na^+^:K^+^:Cl^−^ cotransporter NKCC1 is reorganized in KCC4 due to TM-TM displacement and loss of specific coordinating side chains, explaining Na^+^-independence in KCCs. A second Cl^−^ site in NKCC1 is unoccupied in KCC4, despite being structurally conserved, suggesting coupled binding of Na^+^:Cl^−^ to these sites in NKCC1 and their coincident loss in KCCs.

## Methods

### Cloning and Protein Expression

Cloning, expression, and purification were performed similarly to that described for LRRC8A^26^. The sequence for KCC4 from *Mus musculus* was codon optimized for *Spodoptera frugiperda* and synthesized (Gen9, Cambridge, MA). The sequence was cloned into a custom vector based on the pACEBAC1 backbone (MultiBac; Geneva Biotech, Geneva, Switzerland) with an added C-terminal PreScission protease (PPX) cleavage site, linker sequence, superfolder GFP (sfGFP), and 7xHis tag, generating a construct for expression of mmKCC4-SNS-LEVLFQGP-SRGGSGAAAGSGSGS-sfGFP-GSS-7xHis. Mutations were introduced using standard PCR techniques with partially overlapping primers. MultiBac cells were used to generate bacmids according to manufacturer’s instructions. *Spodoptera frugiperda* (Sf9) cells were cultured in ESF 921 medium (Expression Systems, Davis, CA) and P1 virus was generated from cells transfected with Escort IV Transfection Reagent (Sigma, Carlsbad, CA) according to manufacturer’s instructions. P2 virus was generated by infecting cells at 2×10^6^ cells/mL with P1 virus at a MOI ∼ 0.1. Infection monitored by fluorescence of sfGFP-tagged protein and P2 virus was harvested at 72 hours post infection. P3 virus was generated in a similar manner to expand the viral stock. The P3 viral stock was then used to infect 1 L of Sf9 cells at 4×10^6^ cells/mL at a MOI ∼ 2–5. At 60 hours post-infection, cells were harvested by centrifugation at 2500 x g and frozen at −80°C.

### Transporter Assay

The FluxOR-Red Potassium Ion Channel Assay (Thermo Fisher Scientific) was adapted for transport assays in Sf9 insect cells by adjusting the osmolarity of all buffers to 380 mOsm (by addition of sodium methylsulfonate). Cells were infected at a density of 1.5×10^6^ cells/ml and grown in suspension for 60-72 hours for robust KCC4-GFP expression. 100uL of cells at 1×10^6^ cells/ml were plated and allowed to adhere for 1 hour before the assay. Growth media was replaced with 1X Loading Buffer and incubated at 27°C away from light for 1 hr. The FluxOR Red reagent is a non-fluorescent indicator dye which is loaded into cells as a membrane-permeable acetoxymethyl (AM)-ester. The non-fluorescent AM ester of the FluxOR Red reagent is cleaved by endogenous esterases into a fluorogenic Tl+-sensitive indicator. 1X Loading Buffer was subsequently removed and replaced with Dye-free Assay Buffer and FluxOR Background Suppressor. The assay was performed in 96-well, black-walled, clear-bottom plates (Costar). Fluorescence was measured on a Perkin-Elmer Envision Multilabel Plate Reader using bottom read fluorescence and a BODIPY TMR FP filter set (excitation 531 nm and 25 nm bandwidth, emission 595 nm and 60 nm bandwidth). Fluorescence measurements were made every 0.6 seconds for 300 counts. The recordings were baseline corrected by subtracting the average fluorescence from 180 seconds prior to the addition of Basal Potassium Stimulus buffer and time zero is defined as the first data point recorded after the addition of stimulus. Global fits of all data to a one phase association model Y=(Plateau)*(1-e^(-x/τ)^) are displayed with 95% confidence interval bands (Figure 1B). Alternatively, the final 50 counts were averaged as a measure of final fluorescence increase (Fig 1C, 4F).

### Protein purification

Cells from 1 L of culture (∼7-12.5 mL of cell pellet) were thawed in 100 mL of Lysis Buffer (50 mM Tris, 150 mM KCl, 1 mM EDTA, pH 8.0). Protease inhibitors were added to the lysis buffer immediately before use (final concentrations: E64 (1 µM), Pepstatin A (1 µg/mL), Soy Trypsin Inhibitor (10 µg/mL), Benzimidine (1 mM), Aprotinin (1 µg/mL), Leupeptin (1 µg/mL), and PMSF (1 mM)). Benzonase (5 µl) was added after cells thawed. Cells were then lysed by sonication and centrifuged at 150,000 x g for 45 min. The supernatant was discarded and residual nucleic acid was from the top of the membrane pellet by rinsing with DPBS. Membrane pellets were transferred to a glass dounce homogenizer containing Extraction Buffer (50 mM Tris, 150 mM KCl, 1 mM EDTA, 1% w/v n-Dodecyl-β-D-Maltopyranoside (DDM, Anatrace, Maumee, OH), 0.2% w/v Cholesterol Hemisuccinate Tris Salt (CHS, Anatrace), pH 8.0). A 10%/2% w/v solution of DDM/CHS was dissolved and clarified by bath sonication in 200 mM Tris pH 8.0 and subsequently added to buffers at the indicated final concentrations. Membrane pellets were homogenized in Extraction Buffer and this mixture (100 mL final volume) was gently stirred at 4°C for 1 hr. The extraction mixture was centrifuged at 33,000 x g for 45 min. The supernatant, containing solubilized KCC4-sfGFP, was bound to 5 mL of Sepharose resin coupled to anti-GFP nanobody for 1 hour at 4°C. The resin was collected in a column and washed with 20 mL of Buffer 1 (20 mM Tris, 150 mM KCl, 1 mM EDTA, 1% DDM, 0.2% CHS, pH 8.0), 50 mL of Buffer 2 (20 mM Tris, 500 mM KCl, 1 mM EDTA, 1% DDM, 0.2% CHS, pH 8.0), and 20 mL of Buffer 1. Washed resin was resuspended in 6 mL of Buffer 1 with 0.5 mg of PPX and rocked gently in the capped column overnight. Cleaved KCC4 protein was eluted with an additional 25 mL of Buffer 1. The eluted pool was concentrated to ∼500 µl with an Amicon Ultra spin concentrator 100 kDa cutoff (MilliporeSigma, USA) and subjected to size exclusion chromatography using a Superose 6 Increase column (GE Healthcare, Chicago, IL) run in Buffer 3 (20 mM Tris pH 8.0, 150 mM KCl, 1 mM EDTA, 1% DDM, 0.01% CHS) on a NGC system (Bio-Rad, Hercules, CA). Peak fractions containing KCC4 transporter were collected and concentrated.

### Cross linking and mass spectrometry

Fractions corresponding to peaks 1 and 2 from size exclusion chromatography were separately pooled and concentrated to 0.5 mg/mL. Crosslinking was performed by adding 1 uL of glutaraldehyde from 10X stock solutions in water to 10 uL of KCC4 to achieve final glutaraldehyde concentrations of 0.02, 0.01, 0.005, 0.0025, and 0%. Samples were incubated for 30 minutes prior to quenching by addition of 1uL 1M Tris-HCl and analysis by SDS-PAGE on 4 - 12% Tris-glycine gel (BioRad, USA). Deglycosylated samples were pretreated with 1:10 volume purified PNGase at 1 mg/mL (Addgene 114274) for 1h at 4°C prior to the addition of glutaraldehyde.

For mass spectrometry, the band corresponding to purified KCC4 was excised from a 4 - 12% Tris-glycine gel, digested with trypsin in situ, and the resulting peptides extracted and concentrated. Mass spectrometry was performed by the Vincent J. Coates Proteomics/Mass Spectrometry Laboratory at UC Berkeley. A nano LC column was packed in a 100 μm inner diameter glass capillary with an emitter tip. The column consisted of 10 cm of Polaris c18 5 μm packing material (Varian), followed by 4 cm of Partisphere 5 SCX (Whatman). The column was loaded by use of a pressure bomb and washed extensively with buffer A (5% acetonitrile/ 0.02% heptaflurobutyric acid (HBFA)). The column was then directly coupled to an electrospray ionization source mounted on a Thermo-Fisher LTQ XL linear ion trap mass spectrometer. An Agilent 1200 HPLC equipped with a split line so as to deliver a flow rate of 300 nl/min was used for chromatography. Peptides were eluted using a 4-step MudPIT procedure^42^. Buffer A was 5% acetonitrile/ 0.02% heptaflurobutyric acid (HBFA); buffer B was 80% acetonitrile/ 0.02% HBFA. Buffer C was 250 mM ammonium acetate/ 5% acetonitrile/ 0.02% HBFA; buffer D was same as buffer C, but with 500 mM ammonium acetate.

Protein identification was done with IntegratedProteomics Pipeline (IP2, Integrated Proteomics Applications, Inc. San Diego, CA) using ProLuCID/Sequest, DTASelect2 and Census^43–45^. Tandem mass spectra were extracted into ms1 and ms2 files from raw files using RawExtractor^46^. Data was searched against a *Spodoptera frugiperda* protein database with the purified mouse KCC4 sequence added, supplemented with sequences of common contaminants, and concatenated to a to form a decoy database^47^. LTQ data was searched with 3000.0 milli-amu precursor tolerance and the fragment ions were restricted to a 600.0 ppm tolerance. All searches were parallelized and searched on the VJC proteomics cluster. Search space included all half tryptic peptide candidates with no missed cleavage restrictions. Carbamidomethylation (+57.02146) of cysteine was considered a static modification. In order to identify authentic termini, we required one tryptic terminus for each peptide identification. The ProLuCID search results were assembled and filtered using the DTASelect program with a peptide false discovery rate (FDR) of 0.001 for single peptides and a peptide FDR of 0.005 for additional peptides for the same protein. Under such filtering conditions, the estimated false discovery rate was less than 1%.

### Nanodisc reconstitution

Freshly purified and concentrated KCC4 in Buffer 3 was reconstituted into MSP1D1 nanodiscs with a mixture of lipids (DOPE:POPS:POPC at 2:1:1 molar ratio, Avanti, Alabaster, Alabama) at a final molar ratio of KCC4:MSP1D1:lipids of 0.2:1:50. Lipids in chloroform were prepared by mixing, drying under argon, washing with pentane, drying under argon, and placed under vacuum overnight. The dried lipid mixture was rehydrated in Buffer 4 (20 mM Tris, 150 mM KCl, 1 mM EDTA pH 8.0) and clarified by bath sonication. DDM was added to a final concentration of 8 mM and the detergent solubilized lipids were sonicated until clear. Lipids, Buffer 4 containing 8 mM DDM, and KCC4 protein were mixed and incubated at 4°C for 30 min before addition of purified MSP1D1. After addition of MSP1D1, the nanodisc formation solution was 47.5 µM KCC4, 104 µM MSP1D1, 13 mM DOPE:POPS:POPC, and 4 mM DDM in Buffer 4 (final concentrations). After mixing at 4°C for 30 mins, 60 mg of Biobeads SM2 (Bio-Rad, USA) (prepared by sequential washing in methanol, water, and Buffer 4 and weighed damp following bulk liquid removal) were added and the mixture was rotated at 4°C overnight (∼12 hours). Nanodisc-containing supernatant was collected and spun for 10 min at 21,000 x g before loading onto a Superose 6 column in Buffer 4. Peak fractions corresponding to KCC4-MSP1D1 were collected and spin concentrated using a 100 kDa cutoff for grid preparation.

### Grid preparation

The KCC4-MSP1D1 nanodisc sample was concentrated to ∼1 mg/mL and centrifuged at 21,000 x g for 10 min at 4°C prior to grid preparation. A 3 uL drop of protein was applied to a freshly glow discharged Holey Carbon, 400 mesh R 1.2/1.3 gold grid (Quantifoil, Großlöbichau, Germany). A Vitrobot Mark IV (FEI / Thermo Scientific, USA) was utilized for plunge freezing in liquid ethane with the following settings: 4°C, 100% humidity, 1 blot force, 3s blot time, 5s wait time. The KCC4 detergent sample was frozen at 4.5 mg/mL and centrifuged at 21,000 x g for 10 min at 4°C prior to grid preparation. A 3 µL drop of protein was applied to a freshly glow discharged Holey Carbon, 400 mesh R 1.2/1.3 gold grid. A Vitrobot Mark IV (FEI / Thermo Scientific, USA) was utilized for plunge freezing in liquid ethane with the following settings: 4°C, 100% humidity, 1 blot force, 4s blot time, 1s wait time. Grids were clipped in autoloader cartridges for data collection.

### Data collection

KCC4-MSP1D1 grids were transferred to a Talos Arctica cryo-electron microscope (FEI / Thermo Scientific, USA) operated at an acceleration voltage of 200 kV. Images were recorded in an automated fashion with SerialEM^48^ using image shift with a target defocus range of −0.7 ∼ −2.2 µm over 5s as 50 subframes with a K3 direct electron detector (Gatan, USA) in super-resolution mode with a super-resolution pixel size of 0.5685 Å. The electron dose was 9.333 e^−^ / Å^2^ / s (0.9333 e^−^/ Å2/frame) at the detector level and total accumulated dose was 46.665 e-/Å2. KCC4-detergent grids were transferred to a Titan Krios cryo-electron microscope (FEI / Thermo Scientific, USA) operated at an acceleration voltage of 300 kV. Images were recorded in an automated fashion with SerialEM^48^ with a target defocus range of −0.7 to −2.2 µm over 9.6 s as 48 subframes with a K2 direct electron detector (Gatan, USA) in super-resolution mode with a super-resolution pixel size of 0.5746 Å. The electron dose was 6.092 e^−^ / Å^2^ / s (1.2184 e^−^ / Å^2^ / frame) at the detector level and total accumulated dose was 58.4832 e^−^/Å^2^. See also Table 1 for data collection statistics.

### Data processing

The processing pipeline is shown in Figure S5A-C. We used Cryosparc2^49^ for initial model generation and refinement until reconstructions reached 4-5 Å resolution. Bayesian polishing and nanodisc subtraction in Relion 3.0.7^50, 51^ were used to achieve highest resolution reconstructions. While the contribution of disordered or flexible N- and C-terminal regions to alignments is unknown, the remaining 55 kDa asymmetric membrane protein is among the smallest in terms of resolved mass resolved by cryo-EM to date.

A total of 1559 movie stacks were collected, motion-corrected and binned to 1.137 Å/pixel using MotionCor2^52^, and CTF-corrected using Ctffind 4.1.13^53^ (Figure S5A). Micrographs with a Ctffind reported resolution estimate worse than 5 Å were discarded. A small number of particles (∼1000) were picked manually and subjected to two-dimensional classification to generate references for autopicking in Relion. 1,826,000 particles were autopicked and extracted at 2.274 Å/pixel (2x binned) for initial cleanup. Non-particle picks and apparent junk particles were removed by several rounds of two-dimensional class averaging. The remaining 887,132 particles were extracted at 1.137 Å/pixel and imported into Cryosparc. An additional round of 2D classification generated a particle set of 491,111. These particles were the input of an ab initio reconstruction (non-default values: 4 classes, 0.1 class similarity, 4 Å max resolution, per-image optimal scales). 2D classification of particles (160,868) that contributed to the most featured volume resulted in a set of 125,593 particles which were the input of an ab initio reconstruction. Alignments were iteratively improved using non-uniform (NU) refinement (0.89 window inner radius, 120 voxel box size, 10 extra final passes, 10 Å low-pass filter, 0.01 batch epsilon, minimize over per-particle scale, 1-4 Å dynamic mask near, 3-8 Å dynamic mask far, 6-10 Å dynamic mask start resolution, 4-6 Å local processing start resolution). Two separate NU refinement output volumes were input into a heterogeneous refinement job of the 886,528 particle set (forced hard classification, 10 Å initial resolution, 5 final full iterations). The more featured class (538,280 particles) was heterogeneously refined (forced hard classification, 10 Å initial resolution, 5 final full iterations) (Figure S5B). The particles and volume from one output (354,234 particles) were input into the first of three iterative NU refinements (10 extra final passes for the second and third iteration).

Particle positions and angles from the final cryoSPARC2 refinement job were input into Relion (using csparc2relion.py from the UCSF PyEM^59^) and 3D refined to generate a 4.18 Å map (6 Å low-pass filter, 0.9 degrees initial sampling, 0.9 degrees local searches) (Figure S5C). A second 3D refinement following Bayesian particle polishing improved the map and reported resolution (4.01 Å) (6 Å low-pass filter, 0.9 degrees initial sampling, 0.9 degrees local searches). CTF refinement with beam tilt group estimation and per-particle defocus was performed, although subsequent 3D refinement did not markedly improve the map. Particle subtraction was performed to remove the contribution of the nanodisc density from alignments and subsequent 3D refinement markedly improved the map (reported resolution 3.86 Å or 3.72 Å after postprocessing) (6 Å low-pass filter, 0.9 degrees initial sampling, 0.9 degrees local searches). A final improvement in map quality and reported resolution and was obtained by removing poor particles with a 3D classification job (2 classes, 10 Å initial low-pass filter, 16 tau fudge, no angular sampling). The final particle set (110,143) was subjected to 3D refinement to generate a final map at 3.72 Å resolution (3.65 Å after postprocessing) (6 Å low-pass filter, 0.9 degrees initial sampling, 0.9 degrees local searches). Particle distribution and local resolution was calculated using Relion (Figure S6A,B). FSCs reported in Figure S6C were calculated using Phenix.mtriage.

### Modeling, refinement, and structure analysis

The final cryo-EM maps were sharpened using Phenix.autosharpen^54^. The structure was modeled de novo in Coot and refined in real space using Phenix.real_space_refine with Ramachandran and NCS restraints. Validation tools in Phenix, EMRinger^55^, and Molprobity^56^ were used to guide iterative rounds of model adjustment in Coot and refinement in Phenix. Cavity measurements were made with HOLE implemented in Coot^57^. Electrostatic potential was calculated using APBS-PDB2PQR^58^ Pymol plugin. Figures were prepared using PyMOL, Chimera, ChimeraX, Fiji, Prism, Adobe Photoshop, and Adobe Illustrator software.

## Acknowledgements

We thank members of the Dr. E. Park and Brohawn laboratories, in particular C. Hoel, T. Turney, and Dr. B. Li, for feedback and critical reading of the manuscript. We thank Dr. D. Toso, Dr. J. Remis, and P. Tobias at the Berkeley Bay Area Cryo-EM facility and UC Berkeley Talos Arctica facility for assistance with microscope setup and data collection. We thank Dr. L. Kohlstaedt for assistance with mass spectrometry. This work used the Vincent J. Proteomics/Mass Spectrometry Laboratory at UC Berkeley, supported in part by NIH S10 Instrumentation Grant S10RR025622. We thank Dr. P. He and Dr. M. West of the High-Throughput Screening Facility (HTSF) at UC Berkeley for technical assistance. This work was performed in part in the HTSF. SGB is a New York Stem Cell Foundation-Robertson Neuroscience Investigator. This work is supported by a UC Berkeley Chancellor’s Fellowship (MSR), a NIGMS postdoctoral fellowship F32GM128263 (DMK), the New York Stem Cell Foundation (SGB), NIGMS grant DP2GM123496-01 (SGB), a McKnight Foundation Scholar Award (SGB), and a Klingenstein-Simons Foundation Fellowship Award (SGB).

## Data availability

The final map of KCC4 in MSP1D1 nanodiscs has been deposited to the Electron Microscopy Data Bank under accession code EMD-20807. Atomic coordinates have been deposited in the PDB under ID 6UKN. Original KCC4 in MSP1D1 nanodiscs micrograph movies have been deposited to EMPIAR.

---

**Supplemental Figure 1.**
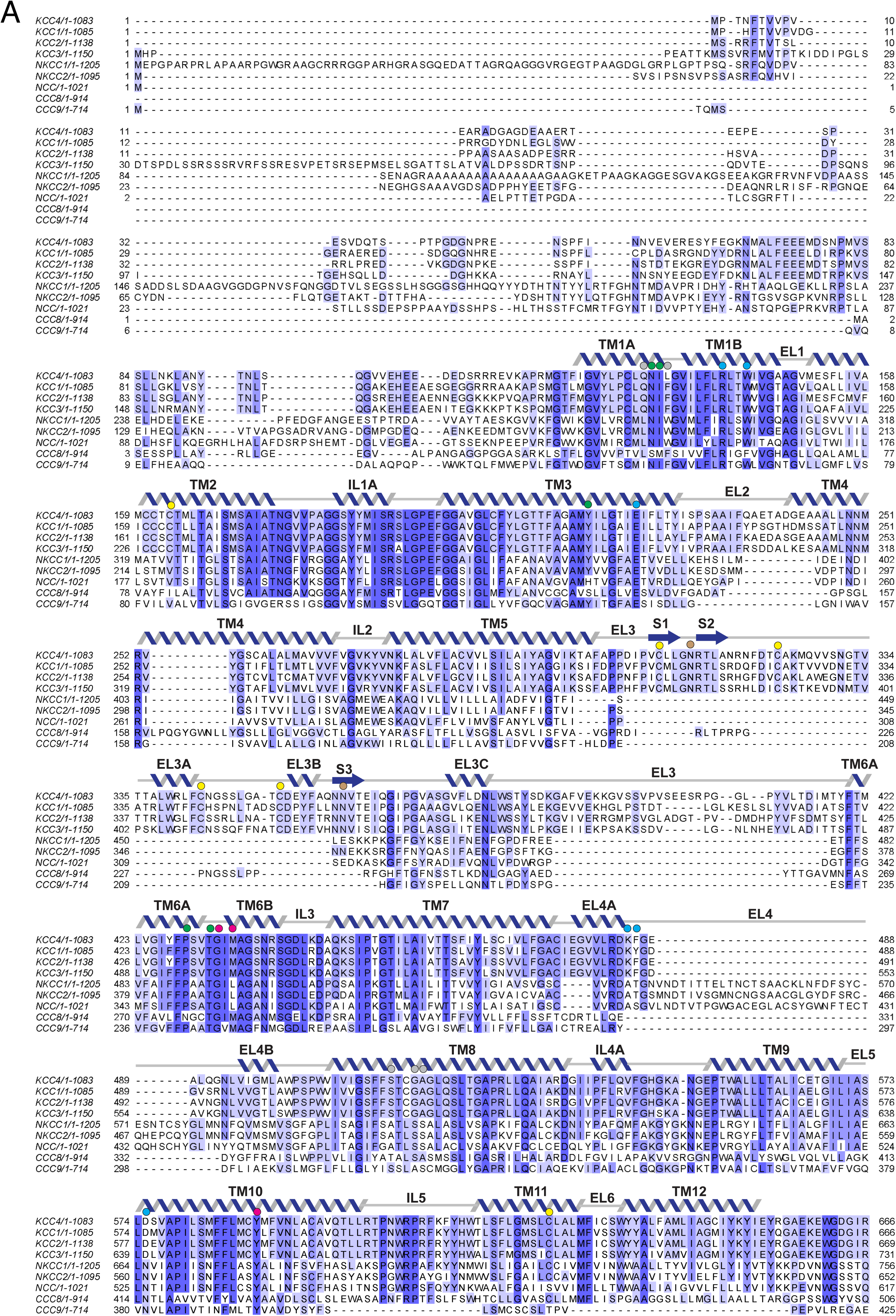
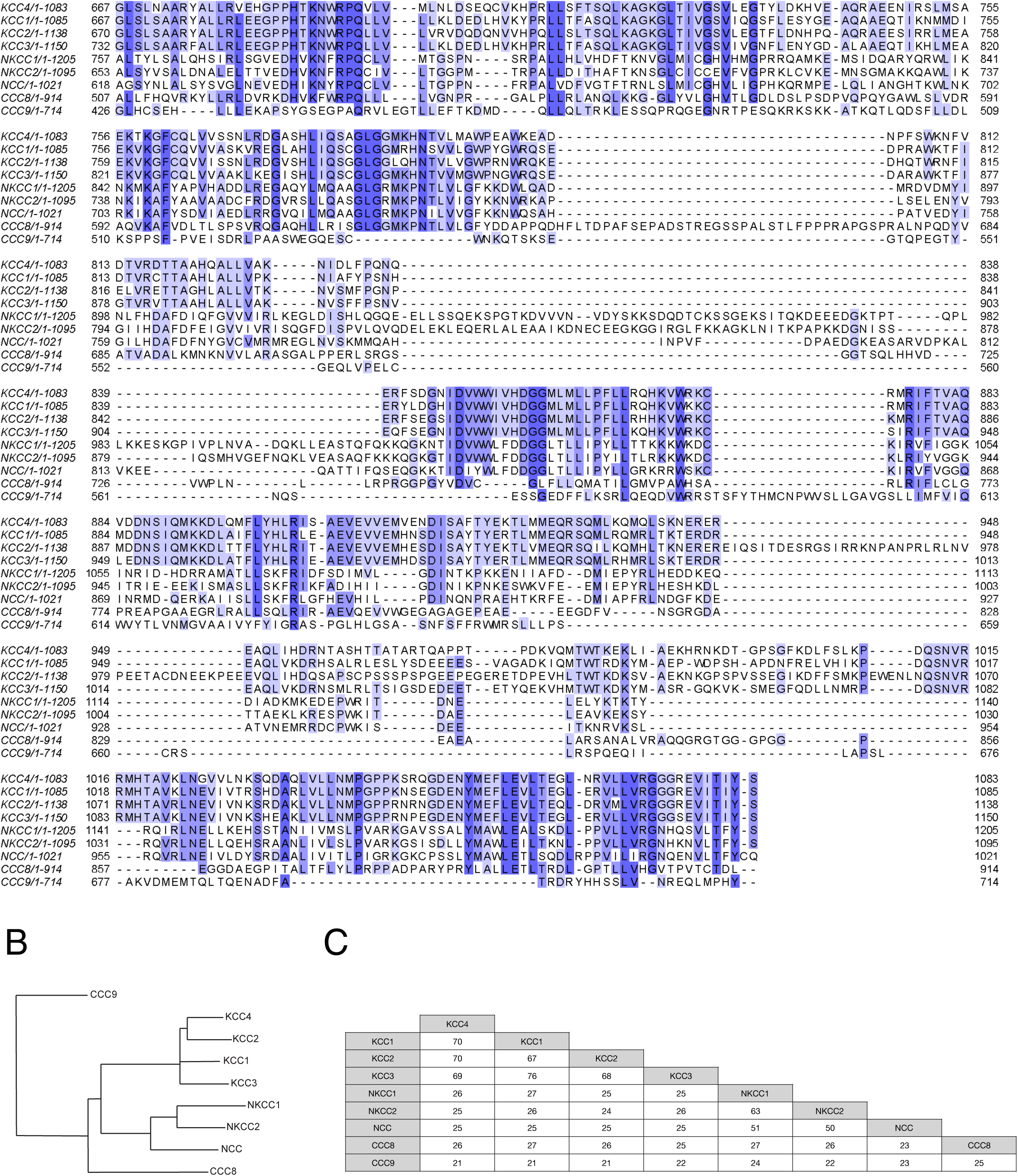
CCC family sequence alignment. (A) Sequence alignment of mouse CCC family transporters. Sequence numbering is indicated at the left and right of each line. Secondary structure for KCC4 is indicated above the sequence. Residues discussed in the text are indicated with circles above the sequence: green, K+ coordinating residues; violet, Cl− coordinating residues; gray, Na+ coordinating residues in NKCC1; yellow, disulfide bond forming residues; brown, glycosylated residues; and cyan, extracellular gate forming residues. (B) Cladogram of CCC family made using sequences from mouse. (C) Percentage sequence identity between members of the mouse CCC family.

**Supplemental Figure 2.**
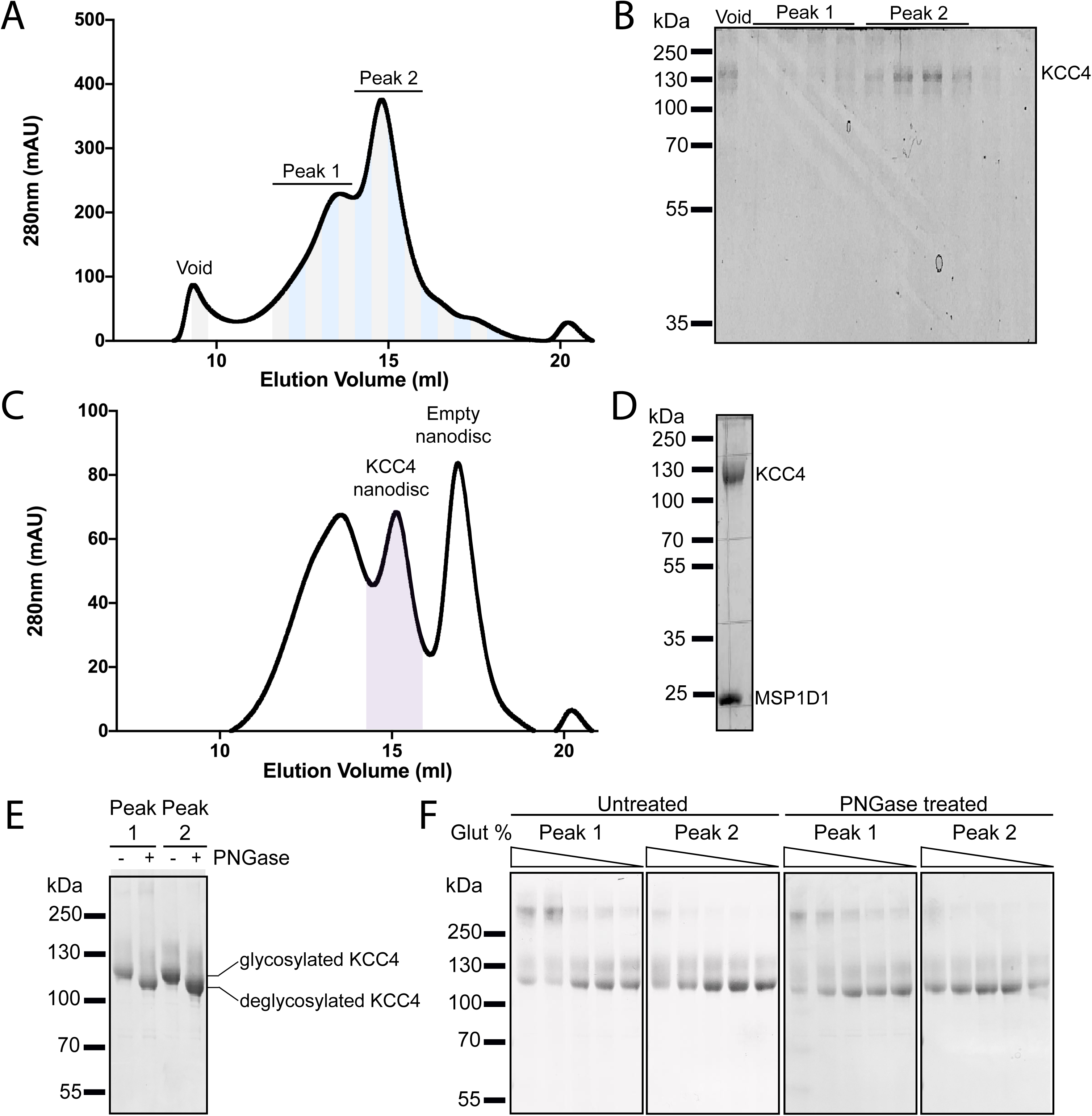
Purification and reconstitution of mouse KCC4. (A) Representative chromatogram from a Superose 6 gel filtration of KCC4 purified in DDM/CHS. (B) Coomassie-stained SDS-PAGE fractions (indicated by alternating gray and blue bands in (A)) with band corresponding to KCC4 labeled. KCC4 is present in two species: an earlier eluting, broader peak 1 and a later eluting, sharper peak 2. Fractions in the later eluting peak correspond to monodisperse KCC4 and were used for structure determination. (C) Representative chromatogram from Superose 6 gel filtration of KCC4 reconstituted in MSP1D1 lipid nanodiscs. (D) Coomassie-stained SDS-PAGE of final pooled KCC4-MSP1D1 nanodisc sample (indicated by purple shading in (C)). (E) Coomassie-stained SDS-PAGE of peak 1 and peak 2 pools (as indicated in (A,B)) before (−) and after (+) treatment with PNGase. (F) Coomassie-stained SDS-PAGE of KCC4 crosslinked with different concentrations of glutaraldehyde (bands within each gel correspond to 0.02, 0.01, 0.005, 0.0025, and 0% glutaraldehyde, respectively). Peak 1 and peak 2 samples were pooled as indicated in (A,B), concentrated, and crosslinked (left pair of gels) or deglycosylated with PNGase prior to crosslinking (right pair of gels).

**Supplemental Figure 3.**
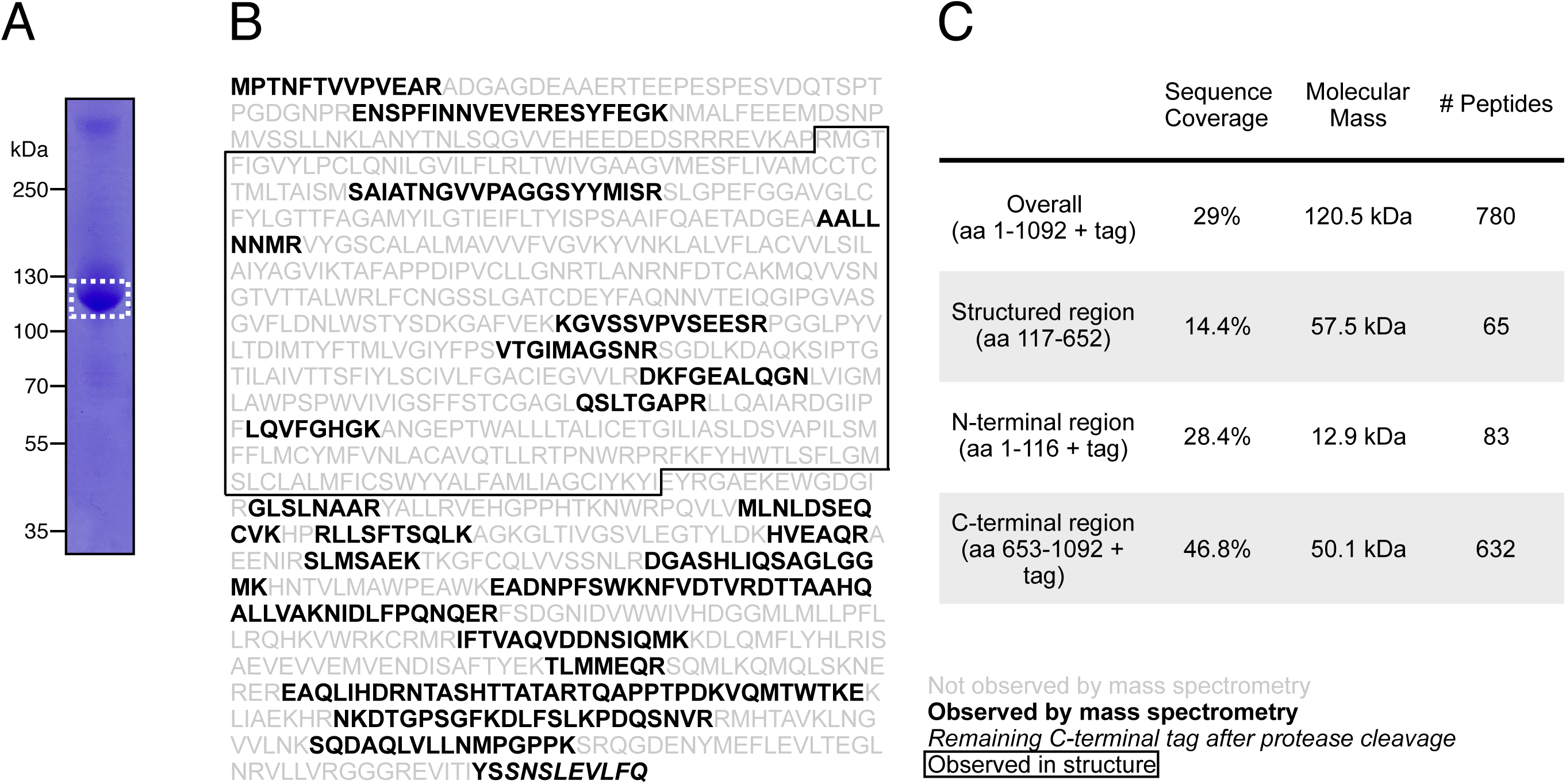
Mass spectrometry of purified KCC4. (A) Coomassie-stained SDS-PAGE of purified mouse KCC4 sample used for mass spectrometry. The band indicated by dashed lines was excised for analysis. (B) Identified peptides (bold) are indicated on the purified KCC4 sequence. Boxed region indicates boundaries of KCC4 observed by cryo-EM. (C) Summary of mass spectrometry data.

**Supplemental Figure 4.**
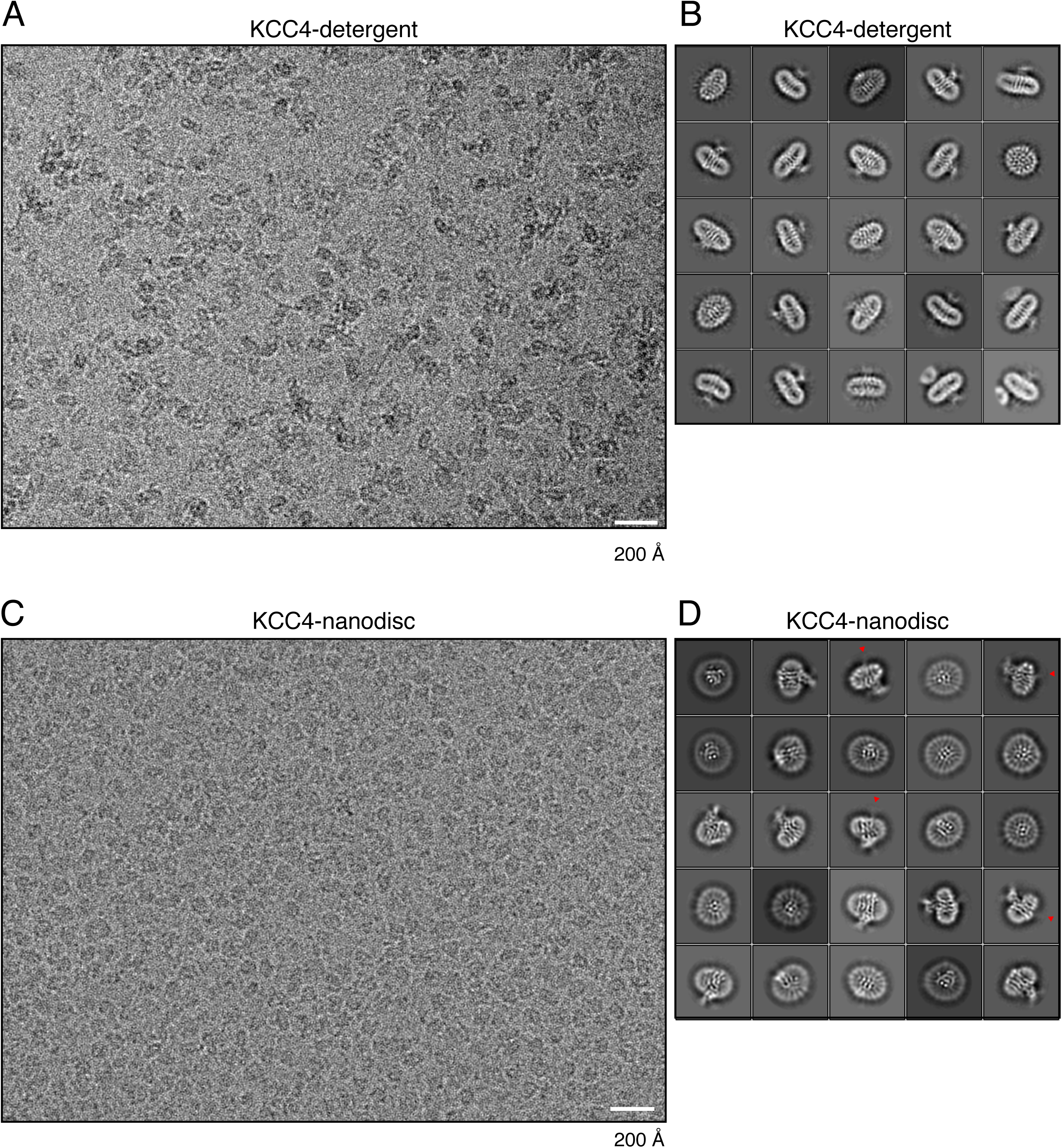
Example micrograph and 2D class averages. (A) Representative micrograph and (B) selected class averages of KCC4 in detergent (DDM/CHS) micelles. (C) Representative micrograph and (D) selected class averages of KCC4 in MSP1D1 lipid nanodiscs. Red arrowheads in (D) point to blurred density features on the intracellular side of the membrane consistent with flexible N-terminal regions or CTDs.

**Supplemental Figure 5.**
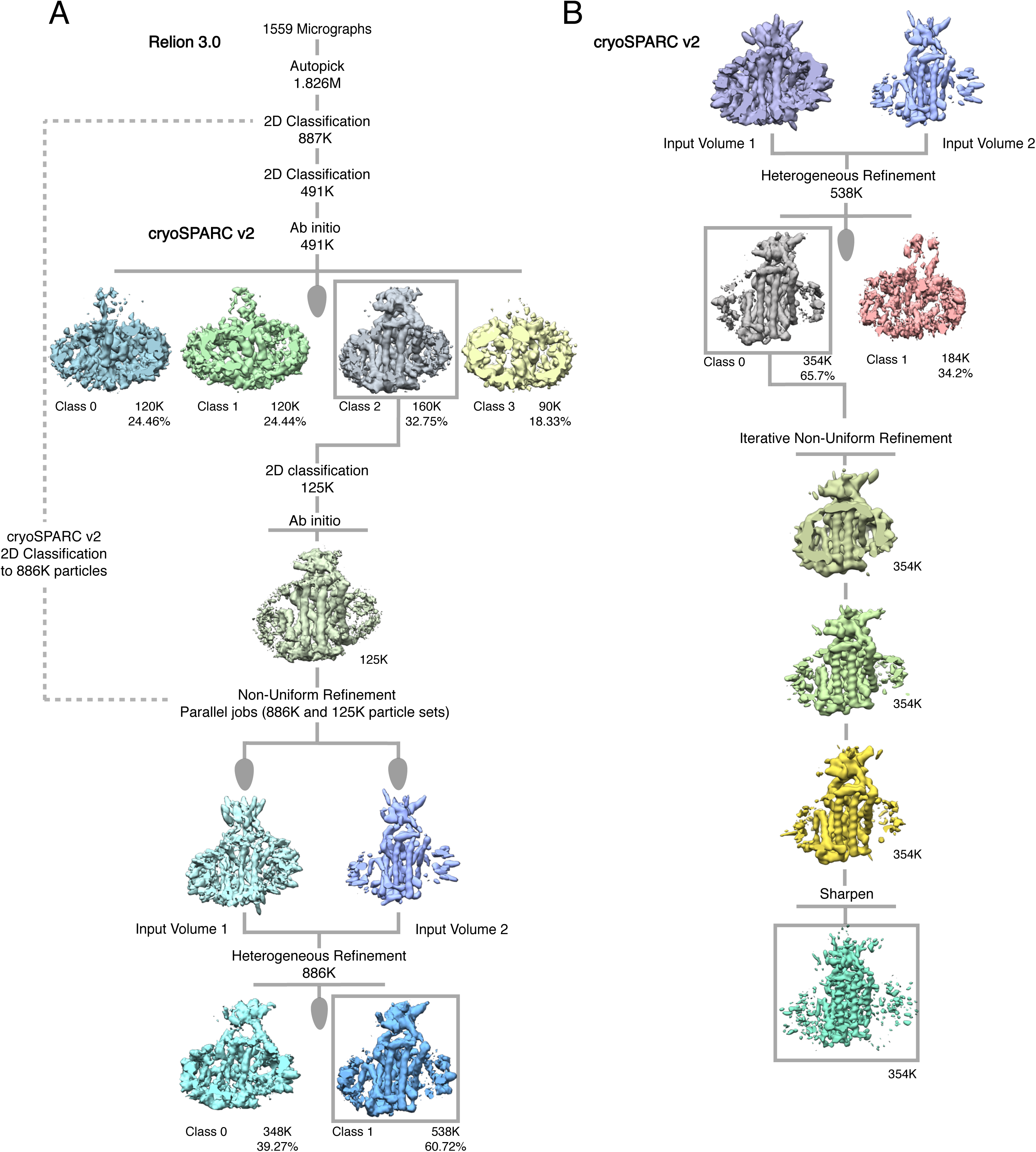
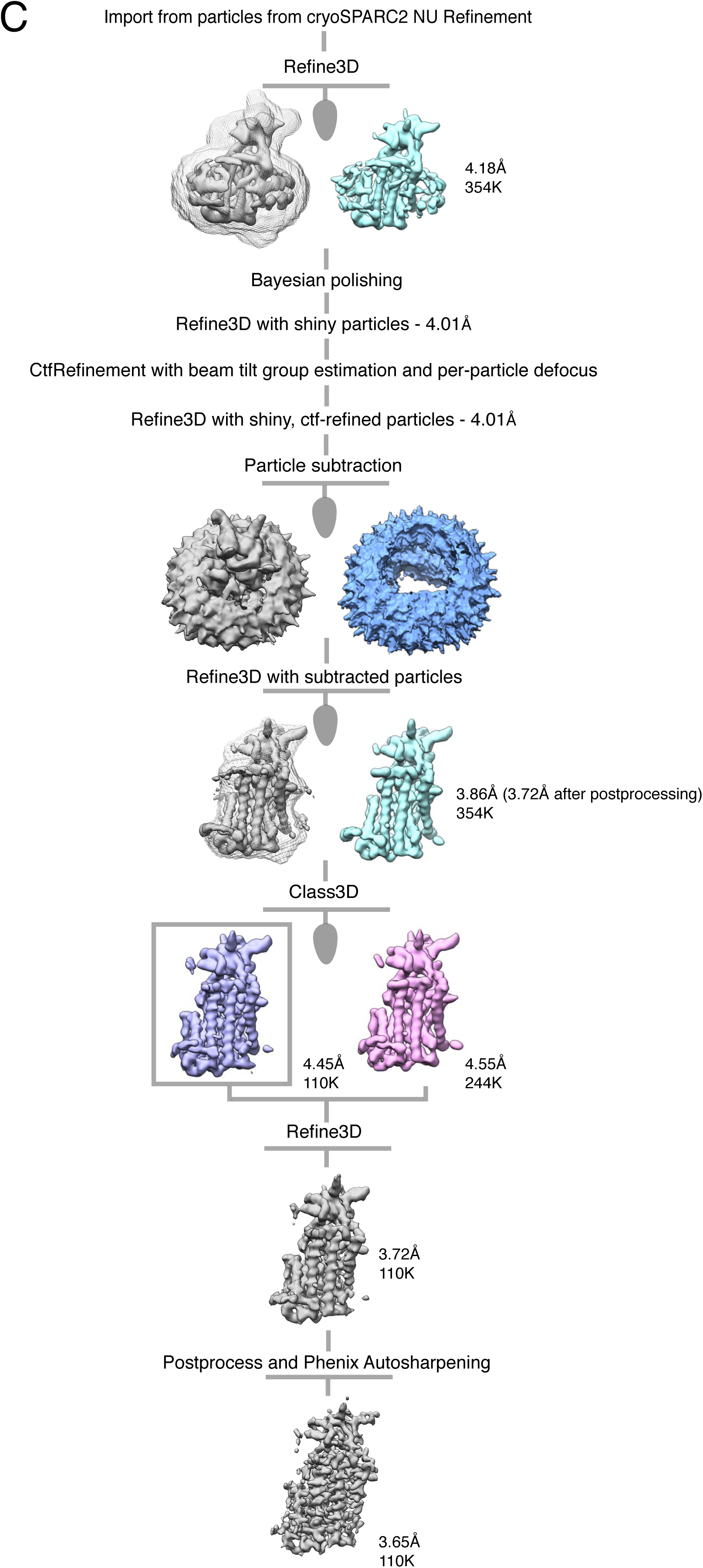
Cryo-EM processing pipeline for KCC4 in MSP1D1 nanodiscs. (A,B) Initial stages of cryo-EM data processing in Relion and cryoSPARC2. (C) Final stages of cryo-EM data processing in Relion. See Methods for details

**Supplemental Figure 6.**
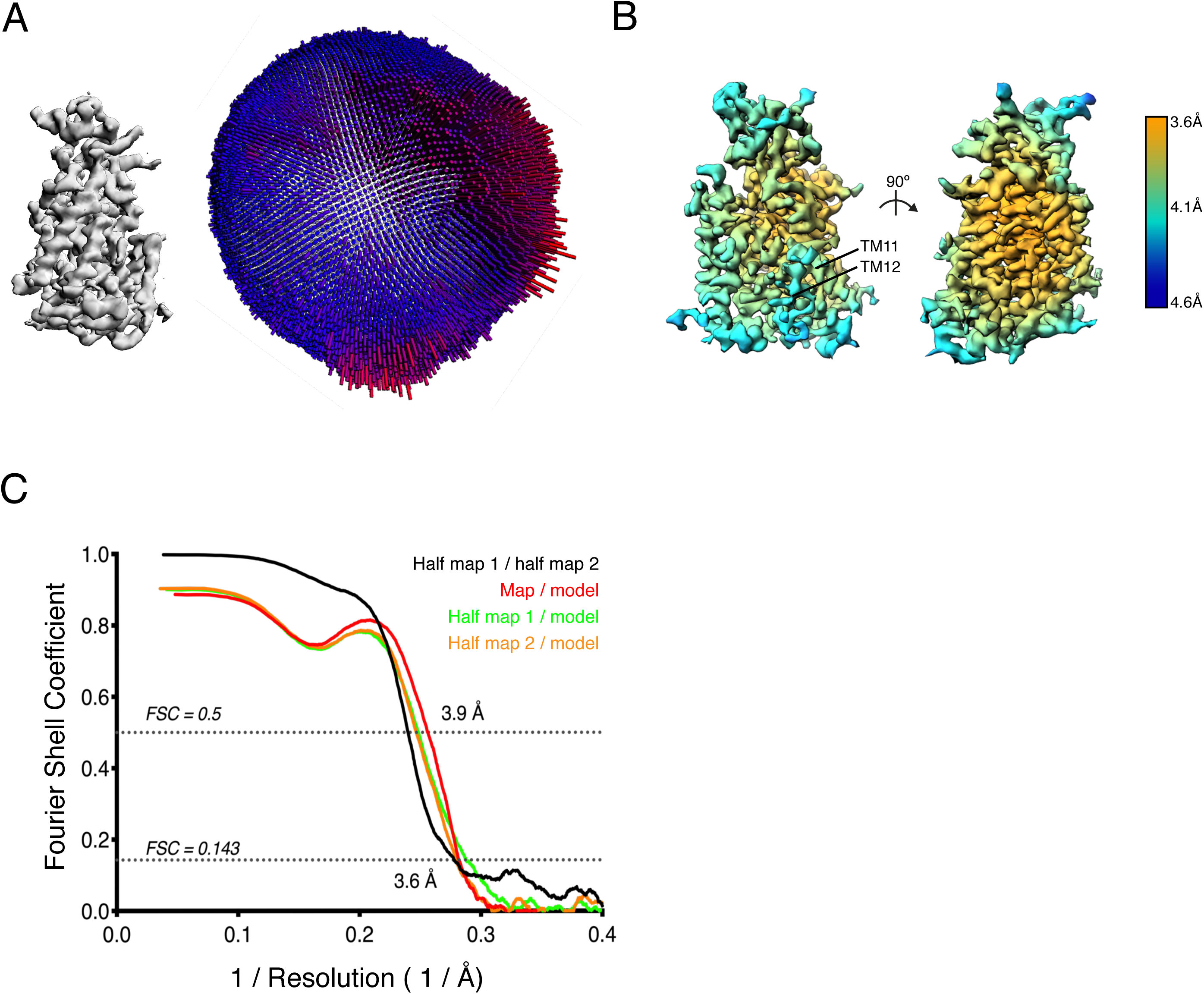
Cryo-EM validation. (A) Angular distribution of particles used in final refinement with final map for reference. (B) Local resolution estimated in Relion colored as indicated on the final map. (Right) a view from the membrane plane showing relatively weaker local resolution in TM11-12 and (left) a view rotated 90°. (C) Fourier Shell Correlation (FSC) relationships between (black) the two unfiltered half-maps from refinement and used for calculating overall resolution at 0.143, (red) final map versus model, (orange) half-map one versus model, and (green) half-map two versus model.

**Supplemental Figure 7.**
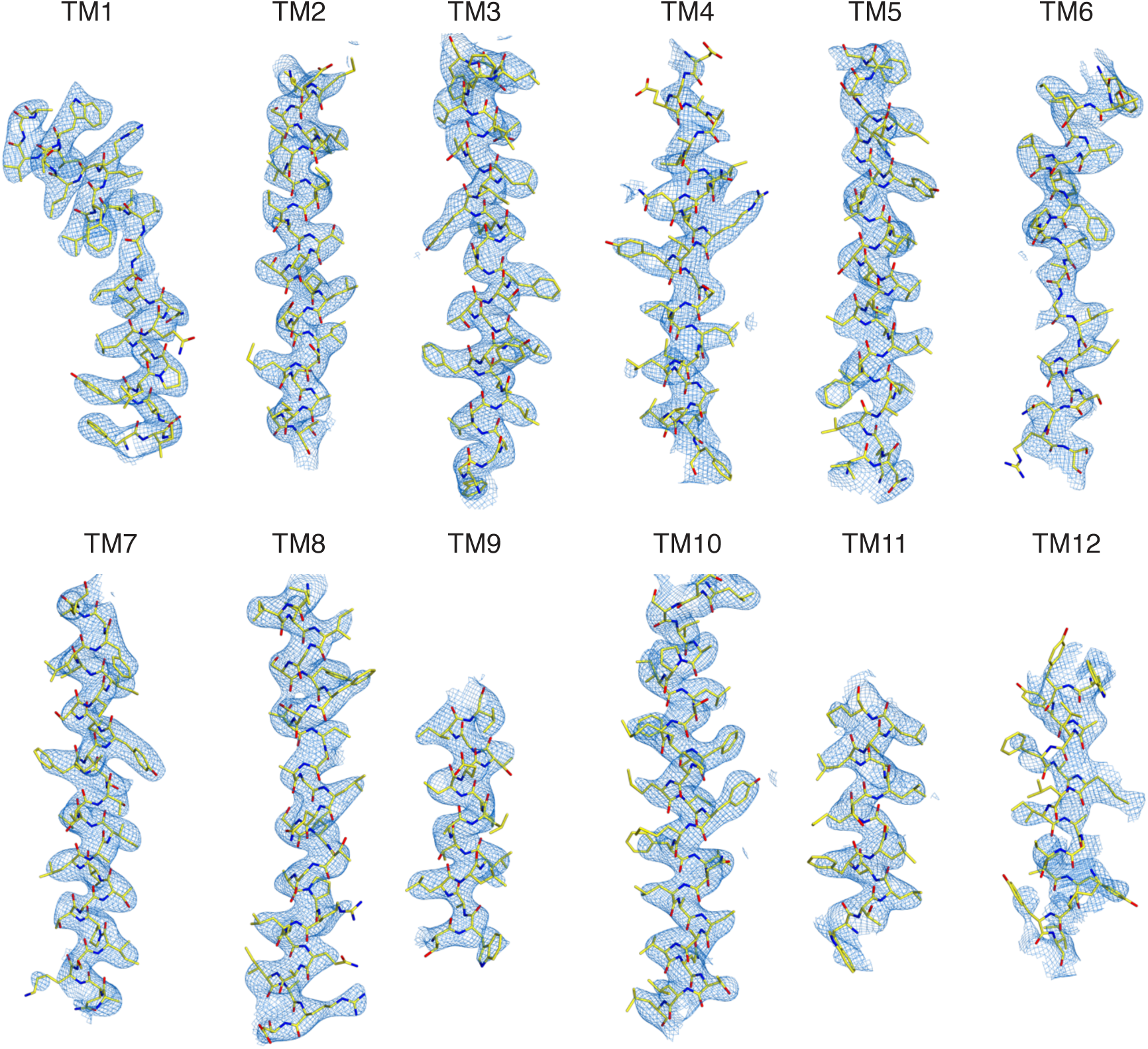
Representative regions of cryo-EM map. Cryo-EM density is shown carved around each transmembrane helix with the atomic model of KCC4 drawn as sticks.

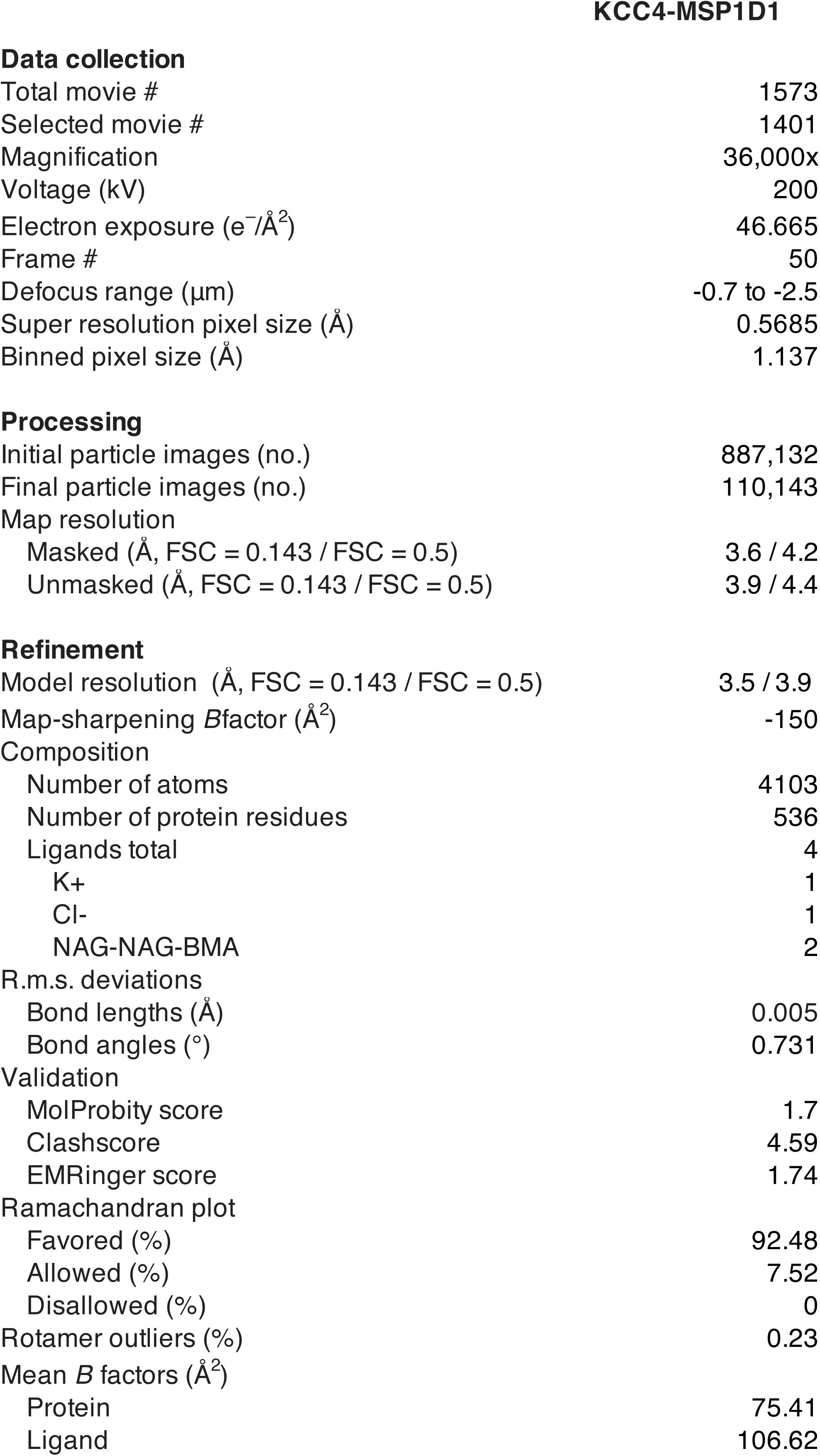

Data collection
|  |  |
| --- | --- |
| Total movie # | 1573 |
| Selected movie # | 1401 |
| Magnification | 36,000x |
| Voltage (kV) | 200 |
| Electron exposure (e<sup>-</sup>/Å<sup>2</sup>) | 46.665 |
| Frame # | 50 |
| Defocus range (μm) | -0.7 to -2.5 |
| Super resolution pixel size (Å) | 0.5685 |
| Binned pixel size (Å) | 1.137 |
| Processing |  |
| Initial particle images (no.) | 887,132 |
| Final particle images (no.) | 110,143 |
| Map resolution |  |
| Masked (Å, FSC = 0.143 / FSC = 0.5) | 3.6 / 4.2 |
| Unmasked (Å, FSC = 0.143 / FSC = 0.5) | 3.9 / 4.4 |
| Refinement |  |
| Model resolution (Å, FSC = 0.143 / FSC = 0.5) | 3.5 / 3.9 |
| Map-sharpening *B*factor (Å<sup>2</sup>) | -150 |
| Composition |  |
| Number of atoms | 4103 |
| Number of protein residues | 536 |
| Ligands total | 4 |
| K+ | 1 |
| Cl- | 1 |
| NAG-NAG-BMA | 2 |
| R.m.s. deviations |  |
| Bond lengths (Å) | 0.005 |
| Bond angles (°) | 0.731 |
| Validation |  |
| MolProbity score | 1.7 |
| Clashscore | 4.59 |
| EMRinger score | 1.74 |
| Ramachandran plot |  |
| Favored (%) | 92.48 |
| Allowed (%) | 7.52 |
| Disallowed (%) | 0 |
| Rotamer outliers (%) | 0.23 |
| Mean *B* factors (Å<sup>2</sup>) |  |
| Protein | 75.41 |
| Ligand | 106.62 |

